# Secretion of small extracellular vesicles is required for commissural axon guidance at a choice point

**DOI:** 10.64898/2026.09.09.750418

**Authors:** Alexandre Dumoulin, Joy M.R. Wyss, Esther T. Stoeckli

## Abstract

The formation of neural circuits relies on precise axonal pathfinding mediated by guidance molecules and their receptors. While the expression of specific guidance molecules by different cell types in the nervous system is well characterized, the mechanisms underlying their secretion and delivery to growing axons remain relatively unexplored. Small extracellular vesicles (sEVs), including exosomes, have emerged as candidate mediators of intercellular communication during nervous system development, with reported roles in neuronal differentiation, synaptogenesis, and axonal outgrowth/regeneration. Whether sEV secretion contributes to axon guidance remains unknown.

Here, we show that axons of dorsal interneurons depend on sEV secretion from floor-plate cells to navigate their intermediate target, the floor plate of the developing chicken spinal cord. Blocking biogenesis of exosomes, a subtype of sEVs, via silencing Rab27a either pharmacologically *ex vivo* or genetically *in vivo*, impairs axon guidance at the floor plate, mainly by interfering with post-crossing navigation. These findings are reproduced after blocking CD63-positive sEV secretion by preventing their formation in multivesicular bodies due to inhibition of VPS4A function. In both cases, axon guidance at the floor plate is perturbed, as axons fail to turn rostral after midline crossing. Taken together, our results show that sEV secretion is required for guidance decisions at a choice point.

## INTRODUCTION

The formation of functional neural circuits relies on the precise synaptic connection between neurons. To reach their target and form synapses, axons have to be accurately guided. More than 30 years of fundamental research has shed light on the roles played by guidance molecules in neural circuit development across many animal models (Dumoulin and Stoeckli, 2023). Initially, the field mainly focused on the identification of guidance molecules, their receptors and downstream-signaling targets (Stoeckli, 2018; Sullivan and Bashaw, 2023). At present, understanding the regulation of axon guidance from a temporal perspective has become the major question. We and others have focused on molecular mechanisms explaining precisely regulated expression of guidance cues and receptors due to changes in transcription (Bourikas et al., 2005; Wilson and Stoeckli, 2013), translation (Baudet et al., 2012; Bellon et al., 2017) or protein stabilization (Gorla et al., 2022; Nawabi et al., 2010). Whereas some guidance molecules are membrane-bound, like Ephrins (Kania and Klein, 2016), other molecules are secreted, like Netrins (Boyer and Gupton, 2018), class-3 Semaphorins (Pasterkamp, 2012), or Sonic hedgehog (Shh) (Zuñiga and Stoeckli, 2017). Still, little is known about how guidance molecules are secreted and delivered to axons.

Small extracellular vesicles (sEVs), in particular exosomes, have been shown to play major roles in cell-cell communication in various systems, such as in cancer progression, in the immune and the nervous system (Buzas, 2023; Filannino et al., 2024; Kalluri and McAndrews, 2023). These membrane-bound nanoparticles (∼50–150 nm in diameter for exosomes) may contain lipids, RNAs, adhesion molecules and signaling molecules (Colombo et al., 2014). They can be secreted in a dynamic fashion and target particular cells by specific recognition/interaction with target cell plasma membrane (Ripoll et al., 2026). Interestingly, multiple studies using mass spectrometry analysis of proteins associated with sEVs identified known axon guidance molecules, such as EphB2, L1CAM, Netrin-1, Plexin A2, Shh, Slit2, or Wnt5a (Chitti et al., 2024; Ditte et al., 2022; Forero et al., 2024; Gong et al., 2016; Sharma et al., 2019; Vyas et al., 2014). This, together with the fact that purified sEV fractions from supernatant of cultured cells showed effects on neurite outgrowth in cultured neurons (Jin et al., 2023; Lai and Breakefield, 2012; Lopez-Verrilli et al., 2016), suggested that these membrane-bound nanoparticles might also play a role during neural circuit formation, particularly in axon guidance (Gong et al., 2016; Lai and Breakefield, 2012; Liu and Teng, 2025; Sharma et al., 2019). However, to the best of our knowledge, no study to date has investigated the possibility that small extracellular vesicles may be important for axon guidance *in vitro* or *in vivo*. Here, using dI1 commissural axons in the developing chicken spinal cord as a model, we show that cells of their intermediate target, the floor plate, secrete small extracellular vesicles, most likely exosomes, while axons are crossing the ventral midline. Impairing exosome biogenesis at two distinct levels, by either silencing Rab27a or by blocking VPS4A in floor-plate cells *in vivo,* causes major axon guidance defects at the contralateral side of the floor plate. Taken together, our results support that small extracellular vesicle or exosome secretion is required for commissural axons guidance at a choice point.

## RESULTS AND DISCUSSION

### Floor-plate cells secrete small extracellular vesicles during commissural axon guidance at the ventral midline

The dorsal-most population of commissural neurons of the developing spinal cord (dI1 subpopulation) is one of the most accessible and best studied models for investigating fundamental molecular mechanisms of axon guidance (Comer et al., 2019; Stoeckli, 2018). The dI1 axons project towards the floor plate (FP) at the ventral midline of the spinal cord and turn rostrally at the contralateral border in a very stereotypical manner between (Hamburger and Hamilton stage; Hamburger and Hamilton, 1992) HH22 and HH26 (Fig. 1A). We used this model and our tools to ask whether secretion of small extracellular vesicles (sEVs) from the FP might play a role in dI1 axon guidance (Fig. 1B). First, we expressed 3 different sEV markers fused to a fluorescent protein specifically in the FP, using *in ovo* electroporation at embryonic day E3. We sacrificed and fixed embryos at E4 (HH23) and visualized the fluorescent signal in transverse slices. We saw positive particles next to the floor plate for all 3 sEV markers, CD63-EGFP, CD81-mCherry, and ALIX-mCherry. This clearly demonstrated that sEVs were secreted at the time when dI1 axons crossed the FP (Figure 1C-E’). We then assessed this secretion by performing live imaging of co-cultures of dorsal spinal cord and FP explants. Upon contact of axons with FP cells, small CD63-EGFP-positive particles were released towards the incoming axonal growth cones (arrowheads, Fig. 1F). Leveraging the use of a pH-dependent CD63 fusion protein (Sung et al., 2020) in these co-cultures, we could visualize CD63-positve particles that were indeed secreted into the medium, as they could be seen on the coverslip and on axons (white arrowheads, Fig. 1G-I). Some sEVs were internalized into axons (yellow arrowheads, Fig. 1I), as they were only positive for mScarlet, but not for pHluo-GFP. Moreover, these observations were reproduced in intact spinal cords cultured *ex vivo.* FP-derived sEVs were secreted within the commissure at HH22, the time when the first dI1 axons are crossing the midline (white and yellow arrowheads, Fig. 1J,K).

**Figure 1.**
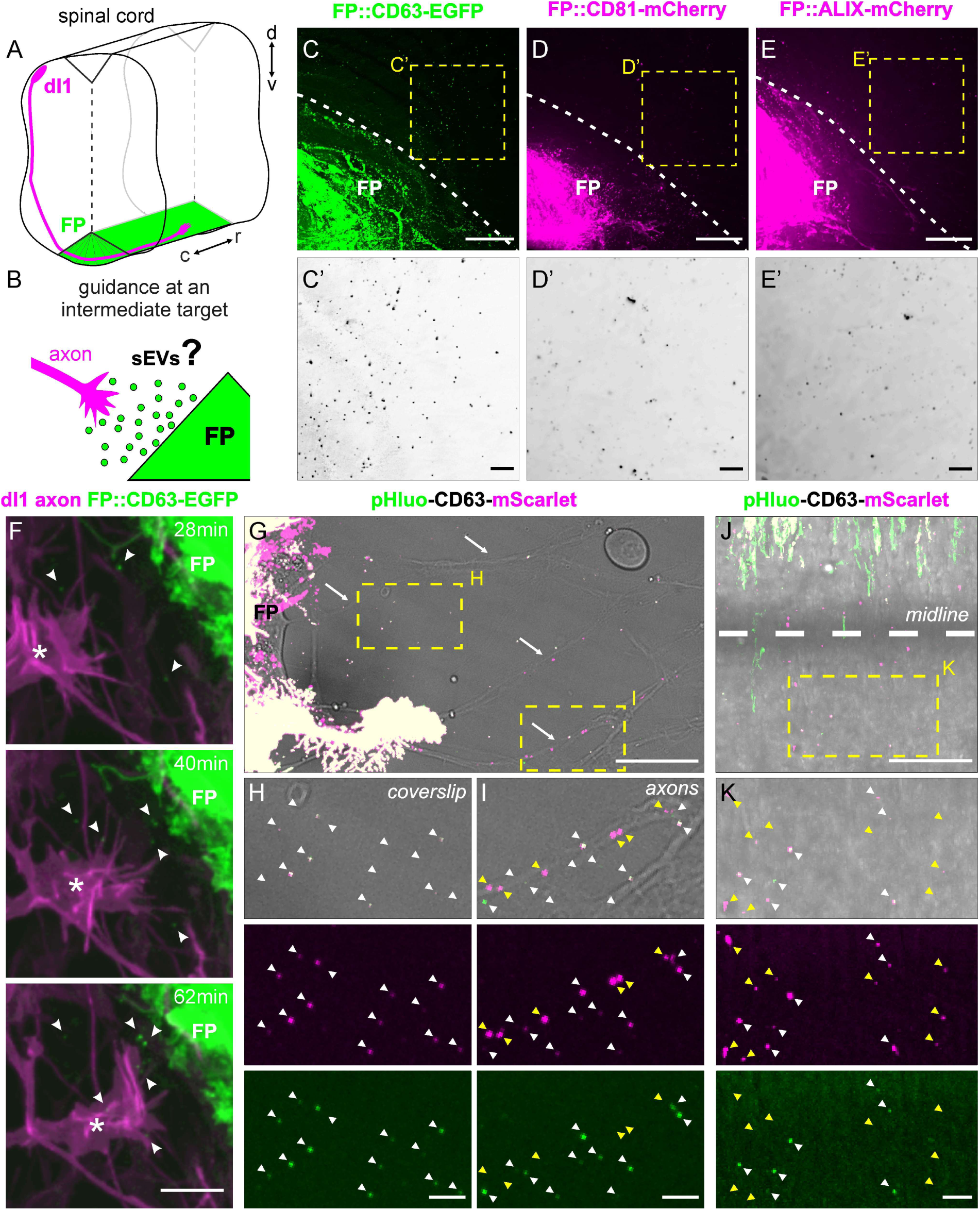
Floor-plate cells secrete small extracellular vesicles during the time window of commissural axon guidance at the ventral midline. (A) Schematic depicting the trajectory of dI1 axons in the developing chicken spinal cord. (B) When dI1 commissural axons approach, enter, and cross the midline, FP cells may secrete sEVs. (B) However, it is currently unknown whether these sEVs play a role in axon guidance. (C-E) Expression of the sEV markers CD63-EGFP (C,C’), CD81-mCherry (D,D’) and ALIX-mCherry (E,E’) in the FP in vivo showed that those markers were also present next to the FP cells (yellow squares, C’-E’). (F) Live imaging of co-cultures of commissural neurons (magenta) and FP explants (green) demonstrated secretion of CD63-EGFP-positive particles (arrowheads) by FP cells upon contact with a dI1 growth cone (asterisk). (G-I) pHluo-CD63-mScarlet allowed visualization of particle secretion upon contact of axons with the FP either on the coverslip surface (H) or on axons (I; white arrowheads). Upper images show a merge with the brightfield channel. Note that some particles were only mScarlet-positive indicating that they were localized intracellularly, and therefore had to be taken up by the axons (I, yellow arrowheads). (J-K) Ex vivo live imaging of an HH22 intact spinal cord expressing pHluo-CD63-mScarlet in one half of the FP showed secreted sEV on the contralateral side of the commissure (white arrowheads) and some that were most likely internalized (yellow arrowheads). d, dorsal; v, ventral; c, caudal; r, rostral. Scale bars: 25 µm (C-E,G,J), 10 µm (F), 5 µm (H,I,K), 1 µm (C’-E’).

Taken together, our results demonstrate that FP cells secrete sEVs when commissural axons cross the midline. This would be compatible with a role in axon guidance.

### Rab27a function in the floor plate is required for commissural axon guidance

Exosome biogenesis happens in two main steps: First, intraluminal vesicles are formed in multivesicular bodies (MVBs). Then, the MVBs dock to and fuse with the plasma membrane for exosome release (Fig. 2A). As the small Rab GTPase Rab27a is known to play an important role in the latter step (Ostrowski et al., 2010), we investigated its expression during the time window of axon guidance. We found expression of its mRNA transcript ubiquitously in the spinal cord (including the FP) throughout stages HH22 to HH26 (arrows, Fig. 2B,C). *In vivo* expression of human Rab27a-EGFP (hRab27a-EGFP) in the FP showed that it localized to the apical as well as the basal feet of FP cells during midline crossing of dI1 axons at HH23 (Fig. 2D). We analyzed midline crossing of dI1 axons over time using *ex vivo* intact spinal cord cultures while blocking Rab27a with Nexinhib-20, a pharmacological Rab27a blocker (Johnson et al., 2016). Inhibition of Rab27a had a strong impact on dI1 axon guidance, as it prevented most axons from turning rostrally at the contralateral side of the FP (red arrows, Fig. 2E, Movie 1). Normal axonal navigation was significantly decreased (15±9%, mean ± s.d.) compared to control conditions (97±3%) with most of the axons showing stalling at the exit site and caudal turns (Fig. 2F,G).

**Figure 2.**
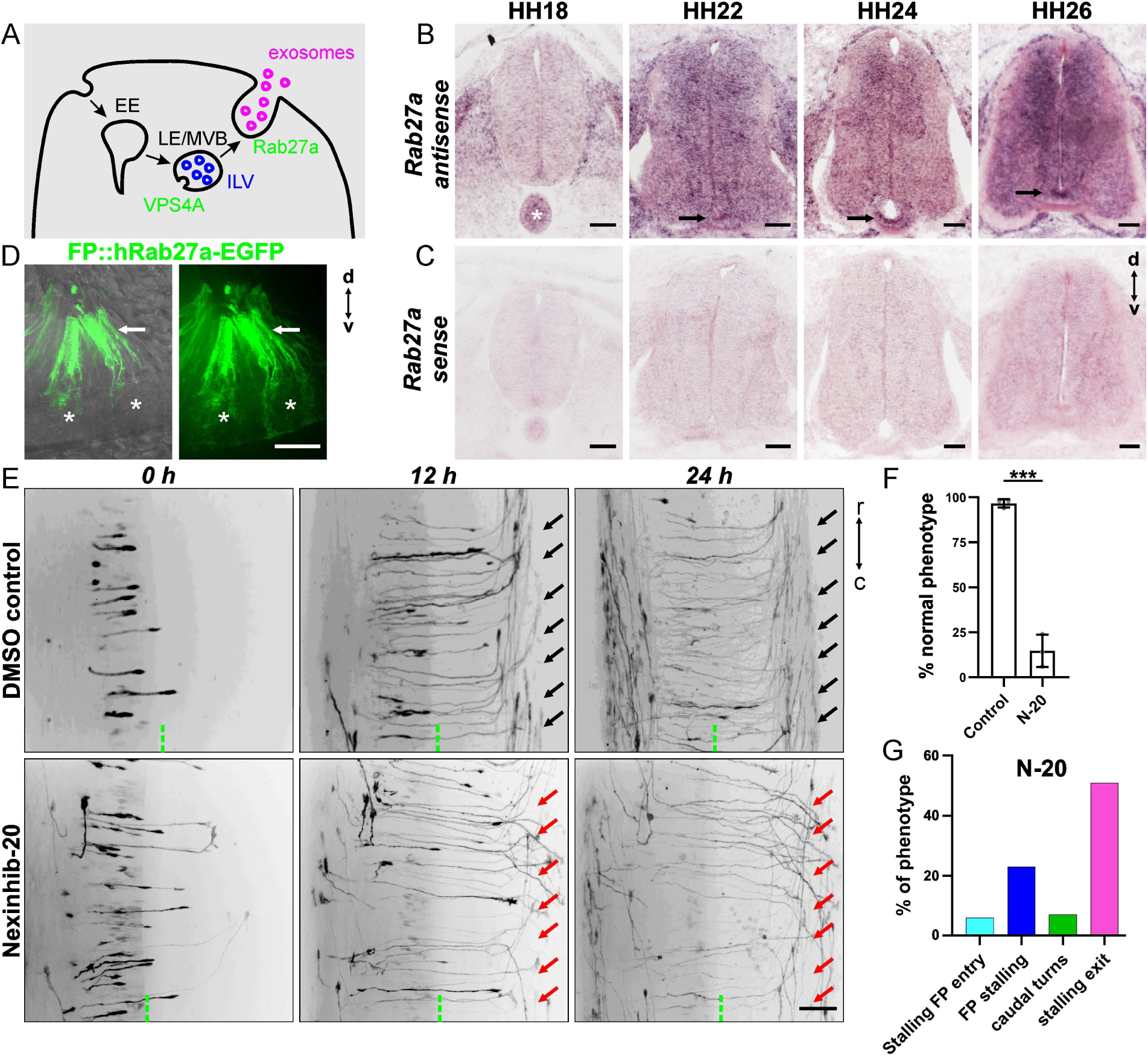
Inhibition of Rab27a in the developing spinal cord impairs commissural axon guidance ex vivo. (A) Cartoon depicting exosome biogenesis. (B) In situ hybridization of Rab27a shows widespread expression of mRNA in the neural tube, including in the FP, during the time window of commissural axon guidance (arrows). (C) Sense probes used as control showed no signal. (D) hRab27a-EGFP expression in the FP in vivo at HH23. Left image shows a merge with the brightfield channel. Arrows point to apical feet, asterisks to basal feet. (E) Inhibition of Rab27a with Nexinhib-20 (N-20) in intact spinal cords ex vivo shows axon guidance defects (red arrows) compared to control (black arrows). (F) Quantification of normal phenotype (rostral turn). N(embryos)=3 (control) and 3 (N-20); n(axons)=273 (control) and 128 (N-20). Error bars represent s.d., ***P<0.001 (two-tailed unpaired t-test). (G) Inhibition of Rab27a mostly induced post-crossing errors, such as stalling at the FP exit site and caudal turning. d, dorsal; v, ventral; c, caudal; r, rostral. Scale bars: 50 µm (C,E), 25 µm (D).

Furthermore, knocking down Rab27a in the floor plate *in vivo* using *in ovo* RNAi confirmed that Rab27a was required for commissural axon guidance, as seen using DiI tracing of dI1 axons in fixed open-book preparations (Fig. 3A-C). The percentage of DiI injection sites with normal axonal trajectories was significantly reduced (49±33%) compared to controls (84±16%). At most sites, axons showed post-crossing defects (asterisks, Fig. 3B,D,E). These aberrant phenotypes could be rescued by expressing human Rab27a-EGFP (hRab27a-EGFP, 74±21% normal DiI injection sites) in FP cells (Fig. 3C-E). In order to assess whether silencing Rab27a had an impact on sEV secretion, we expressed CD63-EGFP and visualized its expression in the presence and absence of dsRab27a. We found that CD63-EGFP protein expression was overall unchanged (6629±2315 versus 7603±4188 A.U. (arbitrary units), Fig. 3F-H). However, quantification of CD63-EGFP-postive particles that were secreted from the FP showed that silencing Rab27a reduced the number of secreted particles (122±20) compared to control (153±26) by about 20% (Fig. I-K).

**Figure 3.**
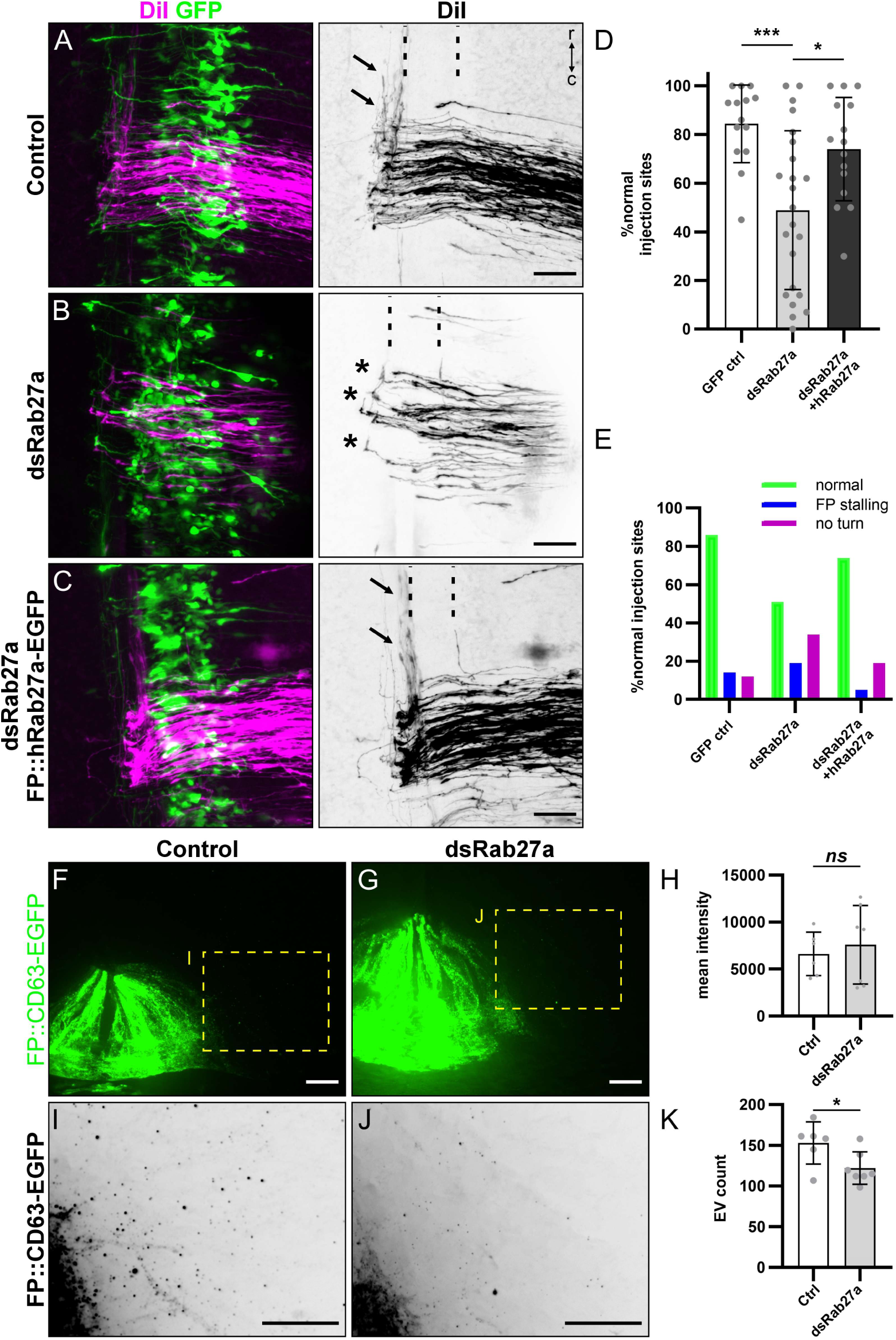
Rab27a plays a non-cell-autonomous role in commissural axon guidance at the midline in vivo. (A) DiI-traced dI1 axons turned normally in GFP-expressing control embryos (A, black arrows), but not in dsRab27a-electroporated embryos (B, asterisks). Co-electroporation of the dsRNA and a plasmid encoding human Rab27a, which is not targeted by the dsRNA, rescued the aberrant phenotype (C, black arrows) (D,E) Quantification of axon guidance phenotypes. N(embryos)=14 (GFP), 23 (dsRab27a), 15 (dsRab27a+hRab27a-EGFP); Error bars represent s.d.; *P<0.05, ***P<0.001 (Brown-Forsythe and Welch ANOVA with Dunnett’s T3 multiple comparisons test). (F,G) Images of HH23 spinal cord transverse slices showing CD63-EGFP expression in the FP with or without transfection of dsRab27a. (H) Quantification of the average mean intensity of the CD63-EGFP signal measured per embryo within the transfected FP. Two-tailed unpaired t test with Welch’s correction. (I,J) Close-up of adjusted images showing secreted CD63-EGFP-positive particles next to the FP. (K) Quantification of the average number of CD63-EGFP-positive particles next to the FP. N(embryos)= 6(Control), 7(dsRab27a). Error bars represent s.d.; *P<0.05, ^ns^P>0.05. two-tailed unpaired t test. r, rostral; c, caudal. Scale bars: 50 µm (A-C), 20 µm (F.G.I,J).

Taken together, our functional analysis of Rab27a, a major regulator of MVB docking to the plasma membrane and exosome secretion, showed its non-cell-autonomous requirement for commissural axon guidance at the midline that correlated with a reduction in FP-derived CD63-EGFP-positive sEV secretion.

### Blocking exosome biogenesis and secretion from the floor plate with cell-specific expression of a dominant-negative form of VPS4A impairs commissural axon guidance

Finally, we wanted to impair the first step of exosome biogenesis, namely the formation of intraluminal vesicles (ILVs) in MVBs by over-expression of a dominant-negative form of VPS4A (VPS4E228Q, VPS4Adn, Fig. 2A). This isoform was shown to impair ILV formation and exosome secretion (Jackson et al., 2017). FP-specific inhibition of VPS4A *in vivo* induced aberrant axon guidance phenotypes at the contralateral border of the FP. Axons were not able to turn rostrally, as in controls, but instead were stalling or turning caudally (Fig. 4A-C). This change of behavior was significantly different (20±17% of normal axonal trajectories) from control (67±31%, Fig. 4D,E). Importantly, overexpression of VPS4dn-EGFP *in vivo* did not yield widespread expression of the protein within FP cells. Its expression was restricted to intracellular compartments and FP cell morphology appeared normal when visualized by the expression of tdTomato in their plasma membrane (Fig. 4F). In order to assess whether blocking VPS4 with this strategy had an impact on sEV secretion, we expressed CD63-EGFP *in vivo* and visualized its expression in the presence of VPS4dn protein. We found that CD63-EGFP protein expression was overall reduced (4349±4658 A.U.) and more restricted to intracellular membrane compartments compared to controls (12760±5389 A.U., Fig. 4G-I). Quantification of CD63-EGFP-postive particles that were secreted from the FP showed that VPS4dn expression drastically reduced their number (60±17) compared to control by about 68% (185±27, Fig. 4J-L). Noteworthy, when we used CD81-mCherry as a reporter for both exosomes and ectosomes in the FP (Mathieu et al., 2021), the expression of VPS4Adn-EGFP reduced also CD81-mCherry expression in the FP compared to controls (1850±1840 A.U. versus 9127±3824 A.U.), similar to the drastic reduction seen for CD63-EGFP (Figure S1A-C). However, it had no impact on the secretion of CD81-mCherry-positive particles from the FP (323±86 versus 344±96, Figure S1D-F). These results demonstrated a correlation between the functional impairment of axon guidance shown above and the reduction of CD63-EGFP-positive sEV secretion by FP cells.

**Figure 4.**
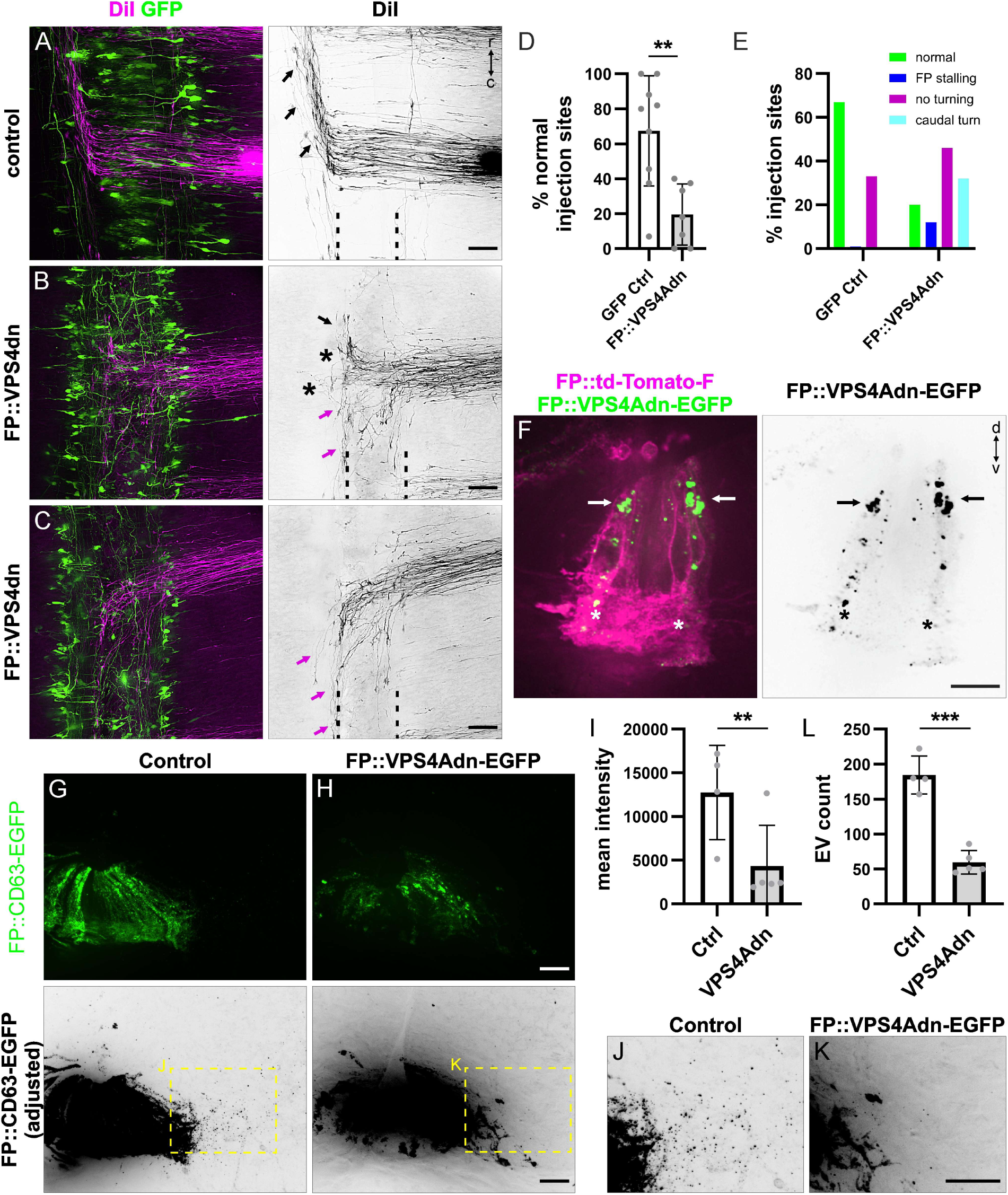
Blockade of floor plate exosome biogenesis and secretion with cell-specific expression of a VPS4A dominant-negative form impairs commissural axon guidance. (A-C) DiI-traced dI1 axons turned normally in GFP-expressing control embryos (black arrows), but not in VPS4dn-electroporated embryos (asterisks and magenta arrows). (D,E) Quantification of axon guidance phenotypes. N(embryos)=9 (GFP), 7 (VPS4Adn); **P<0.01, ***P<0.001 (Unpaired t test with Welch’s correction). (F) Image of a transverse spinal cord slice with td-Tomato-F-expressing FP cells at HH23 showing VPS4dn-EGFP localization in the apical (arrows) and basal feet (asterisks). (G,H) Images of HH23 transverse spinal cord slices showing CD63-EGFP expression in the FP with or without co-expression of VPS4dn. (I) Quantification of the average mean intensity of the CD63-EGFP signal measured per embryo within the transfected FP in vivo. Two-tailed unpaired t test. (J,K) Close-up of adjusted images showing secreted CD63-EGFP-positive particles next to the FP. (L) Quantification of the average CD63-EGFP-positive EV count next to the FP. N(embryos)= 4(Control), 5(VPS4dn). Two-tailed unpaired t test. Error bars represent s.d. d, dorsal; v, ventral; r, rostral; c, caudal. Scale bars: 50 µm (A-C), 20 µm (G-H, J-K), 10 µm (F).

Taken together, our results propose a model that requires exosome secretion from the intermediate target as a means to instruct commissural axons during midline crossing. Our results showed that FP cells secrete sEVs at the time, when axons cross the midline and have to change their behavior by responding to cues secreted by the FP (Fig. 1). Importantly, our functional data demonstrate that sEV secretion from FP cells is not only a coincidence but required for axons to properly navigate their intermediate target. This conclusion is supported by our *in vivo* findings of aberrant axonal navigation after inhibition of sEV secretion either by blocking exosome biogenesis through interference with ILV formation or by interference with MVB docking to the plasma membrane (Figure 2-4). In all these conditions, axons predominantly failed to switch to their post-crossing stage, and thus, did not turn correctly upon floor-plate exit (Fig. S1G).

Our current results do not focus on specific axon guidance molecules that would be transferred by sEVs. However, the fact that the phenotypes obtained after silencing or blocking different components of the exosome biogenesis pathway led to similar post-crossing phenotypes is intriguing. It resembled phenotypes obtained after silencing pathways such as Shh, Wnt or PlexinA2-Sema6B in our system (Andermatt et al., 2014; Bourikas et al., 2005; Domanitskaya et al., 2010). Given that components of these pathways were purified in sEV fractions before (see introduction), it is tempting to speculate that one or more of these pathways might use exosome secretion as a means to transfer a signal to axons crossing the midline. The fact that we identified a drastic reduction of CD63-EGFP-positive sEV secretion, but not CD81-mCherry-positive ones, suggest that the phenotypes reported here were due to the lack of secretion by the FP of a specific subtype of sEVs. Further work will be required to uncover what guidance molecules are secreted by FP-derived sEVs and how they signal to the axonal growth cones to guide them to their targets.

## MATERIALS AND METHODS

### *In ovo* electroporation

*In ovo* electroporation was carried out as previously described in detail (Wilson and Stoeckli, 2012). Solutions containing plasmids encoding fluorescent proteins, tagged proteins and/or dsRNA were injected into the central canal of the embryonic chicken spinal cord at HH18-19. Spinal cord cells were transfected by electroporation using electrodes connected to a BTX ECM830 square-wave electroporator. Five pulses of 25 V, 50 msec duration, 1 s interpulse interval were used for ventral or unilateral electroporation and expression of constructs into the FP for *in vivo*, *ex vivo* and *in vitro* experiments (Table S1). Targeted electroporation was achieved by carefully positioning the electrodes and aligning the dorsal-ventral axis of the neural tube between them using a pair of tweezers to align the embryo by only touching the extra-embryonic membrane. The polarity of the electrodes was set accordingly. Successful targeting was verified by the expression of fluorescent proteins. For successful targeting of the FP cells the Hoxa1 enhancer was used (Dumoulin et al., 2021). In all figures and text ‘FP::xxx’ refers to ‘Hoxa1::xxx’. The list of all plasmids, including their concentrations and origins, is provided in Table S1.

### Plasmids

New plasmids were generated for this study (plasmids marked with an asterisk in Table S1) using the NEBuilder® HiFi DNA Assembly kit (New England Biolabs) to subclone them under a specific promoter/enhancer combination (Hoxa1 enhancer or β-actin promoter)(Wilson and Stoeckli, 2011). The list of all plasmids, including their concentrations for *in ovo* electroporation and their origins, is provided in Table S1.

### *Ex vivo* live imaging of intact spinal cords

*Ex vivo* live imaging of intact spinal cords was carried out as previously described in detail (Dumoulin et al., 2021). In short, intact spinal cords were dissected from HH22 embryos in ice-cold PBS and plated with the ventral midline down for live imaging of axons crossing the midline in a 100-μl drop of 0.5% low-melting agarose diluted in spinal cord medium [MEM with Glutamax (Gibco) supplemented with 4 mg/ml Albumax (Gibco), 1 mM pyruvate (Sigma-Aldrich), 100 units/ml Penicillin and 100 μg/ml Streptomycin (Gibco)] on a 35-mm Ibidi μ-dish with glass bottom (Ibidi, Cat#81158). Once the agarose drop solidified, 200 μl of spinal cord medium were added on top, and samples were incubated at 37°C, with 5% CO_2_ and 95% air, for at least 30 min before live imaging was started. Atmosphere and temperature were controlled with a PeCon cell vivo chamber (PeCon). Images were acquired using an IX83 inverted microscope equipped with a spinning disk unit (CSU-X1 10,000 rpm, Yokogawa), a 40x silicone oil objective (UPLSAPO S 40x/1.25, Olympus), and an Orca-Flash 4.0 camera (Hamamatsu) with the Olympus CellSens Dimension 2.2 software. Stacks (1.4 μm spacing) were acquired at different intervals ranging from 0.5 to 5 min depending on the region of interest that was imaged. For inhibition of Rab27a, a final concentration of 67 µM Nexinhib-20 (SML1919-5MG, Sigma) diluted in DMSO (1:1000 final dilution) was added to the intact spinal cords. For the control condition the equivalent dilution of DMSO (1:1000) was added. Movies of time-lapse recordings were processed and assembled using ImageJ/Fiji (Schindelin et al., 2012).

### Live imaging of explant cultures and co-cultures

Live imaging of explant cultures and co-cultures was conducted as previously described (Dumoulin et al., 2024). The 8-well live imaging plates (µ-Slide 8 Well high, Cat# 80806, ibidi) were coated with 20 μg/ml poly-L-lysine (Sigma-Aldrich, P-12374) and 20 μg/ml laminin (Invitrogen, 23017-015). Pieces of dorsal spinal cord, containing dI1 neurons, were dissected from HH22 open-book preparations and plated together with HH25 FP explants. Spinal cord medium (see above) was used and supplemented with N3 (100 μg/ml transferrin, 10 μg/ml insulin, 20 ng/ml triiodothyronine, 40 nM progesterone, 200 ng/ml corticosterone, 200 μM putrescine, 60 nM sodium selenite; all from Sigma-Aldrich). After plating, explants were incubated at RT for 20 min and then at 37°C, with 5% CO_2_ and 95% air for at least 30 min before live imaging was started. Importantly, FP explants retained their basal-apical polarity in culture. Live imaging conditions and acquisitions were similar to the ones described above for intact spinal cords. For high-resolution temporal acquisition, only single focal planes were acquired using the Z-drift controller to maintain steady focus over time (Olympus). Movies of time-lapse recordings were processed and assembled using ImageJ/Fiji.

### Trunk fixation and transverse slices

HH23-23.5 trunks were fixed for 20 min at RT with 4% paraformaldehyde (PFA) in PBS. Three hundred µm thick slices were cut with a tissue chopper (McIlwain) and mounted immediately on a 24x60 mm cover slip (Menzel-Gläser) in a drop of PBS surrounded by a vacuum grease border with a 24x24 mm cover slip put on top (Menzel-Gläser). Image stacks were taken with an IX83 inverted microscope equipped with a spinning disk unit (CSU-X1 10,000 rpm, Yokogawa), a 40x silicone oil objective (UPLSAPO S 40x/1.25, Olympus), and an Orca-Flash 4.0 camera (Hamamatsu) with the Olympus CellSens Dimension 2.2 software. Images were processed and assembled using ImageJ/Fiji. Note that imaging of sEVs using the reporters used in this study can be challenging if the tissue is over-fixed and not imaged within 1-5 days after fixation. Moreover, the use of any detergent is detrimental to the sEV signal.

### Quantification of secreted sEV *in vivo*

A region of interest consisting of a rectangle of 80x50 µm was taken just next to the FP entry zone where dI1 axons enter it. Contrast and intensity of the signal were adjusted to see sEVs. They were then manually counted using the multi-point tool in Fiji. A minimum of 4 (CD63-EGFP experiments) or 5 slices (CD81-mCherry experiments) per embryo were quantified. For the intensity of the signal, a region of interest was drawn around the transfected FP cells and the mean intensity was measured using Fiji. Both mean intensity values and EV count values were averaged per embryo and statistically tested.

#### Cryosections for *in situ* hybridization

The embryos were sacrificed and dissected at stages HH18 - HH26. The organs were removed and the embryos were fixed in 4% PFA/PBS for 20–45 min, depending on their developmental stage. The embryos were cryoprotected overnight in 25% sucrose in PBS before being embedded in Tissue-Tek O.C.T. Compound (Sakura) and frozen using isopentane on dry ice. Note that all buffers were treated with diethyl pyrocarbonate (DEPC, Applichem). The embryonic trunks were then sectioned transversely into 25-μm-thick sections using a cryostat (LEICA, CM1850) and stored at -20 °C until ISH was performed.

### Preparation of *in situ* hybridization probes

An expressed sequence tag (ChESTs; SourceBioScience, Nottingham, United Kingdom) for Rab27a (ChEST21n27; NM_001395913.1 Gallus gallus RAB27A, member RAS oncogene family (RAB27A), mRNA: 191 – 1331 bp) was used to synthesize probes for in situ hybridization. The ChEST plasmid was linearized with the restriction enzymes NotI and SalI for 4 h at 37 °C. T7 and T3 RNA polymerases (Promega) were used to synthesize the sense and antisense ISH probes, respectively, with a DIG RNA labeling kit (Roche).

#### *In situ* hybridization

The protocol was adapted from (Mauti et al., 2006). Cryosections were air-dried at room temperature before being washed with DEPC-PBS and then with ultrapure H₂O. To inactivate endogenous alkaline phosphatase activity and increase specificity for the target RNA, the sections were immersed in freshly prepared 1% triethanolamine, and 0.25% (v/v) acetic anhydride in ultrapure H₂O. The sections were washed with DEPC-PBS and once with DEPC-2x SSC (0.3 M NaCl, 0.03 M tri-sodium citrate, pH 7). To prevent nonspecific RNA binding, the slides were incubated with 500 μl of prehybridization buffer in a chamber at 56 °C for 4 h (unless otherwise specified, the following steps were performed at 56 °C). For RNA-RNA hybridization, 500 μl of either the antisense or sense probes (1.5 ng/μl), mixed with hybridization buffer, was applied to the sections overnight. The slides were then washed for 5 min each with 5x SSC, 2x SSC, and 0.2x SSC, and for 20 min at 56 °C with 50% formamide in 0.2x SSC. The slides were then transferred to 0.2× SSC at room temperature (RT) for 5 min before being washed twice with detection buffer (0.1 M Tris-base, 0.15 M NaCl, pH 7.5) for 5 minutes each time. To prevent nonspecific antibody binding, 500 μl blocking buffer (3% milk powder in detection buffer) were added to each slide and incubated for 60 min at RT in a humidified chamber. The blocking buffer was replaced with 500 μl of the antibody mixture (1:5000 DIG-AP) per slide and incubated for 120 min at RT. The slides were then washed twice with detection buffer for 15 min each time. To equilibrate the tissue, the slides were treated for 5 min with an alkaline phosphatase (AP) solution (0.1 M Tris-base, 0.1 M NaCl, 50 mM MgCl₂, pH 9.5). To develop the chromogenic signal, 500 μl of the development solution (0.675 mg/ml NBT, 0.24 mg/ml Levamisol, 0.35 mg/ml BCIP in AP buffer) was applied to each slide. The incubation time took approximately 18 h, protected from light. To stop the development, the slides were immersed in Tris-EDTA (TE) buffer (10 mM Tris-base, 1 mM EDTA, pH 8.0) and then washed twice with TE buffer for 10 min each time. The slides were immersed in deionized water before being embedded in Mowiol 4-88 (Carl ROTH; pH 7) as the embedding medium. They were then air-dried overnight and stored at 4 °C.

### ISH imaging acquisition

Images were acquired using an Olympus BX-63 upright microscope equipped with an Olympus DP80 camera. Either a 10x air objective (UPLFLN 2PH 10x / 0.30, Olympus) or a 20x air objective (UPLXAPO 20x / 0.80, Olympus) was used in conjunction with the Olympus CellSens Dimension 2.2 software. To ensure comparability, identical exposure and gain settings were used for imaging the corresponding sense and antisense probes. Fiji (Schindelin et al., 2012) was used for image processing.

### Preparation of dsRNA

For the synthesis of long dsRNA, the Rab27a plasmid was linearized with NotI or Sall, (New England Biolabs, Roche) for the synthesis of sense and antisense single-stranded RNAs (ssRNA) with T7 and T3 RNA polymerase (Promega), respectively. Equal amounts of purified ssRNAs were annealed by allowing the solution to cool slowly to room temperature after heating at 95^°^C for 10 min. Successful double-strand formation was verified by gel electrophoresis. Long dsRNAs were injected and electroporated at a concentration of 500 ng/µl in PBS, together with 25 ng/µl of a reporter plasmid encoding humanized Renilla GFP under the β-actin promoter (β-actin::hrGFPII, termed GFP in results section) and 0.01% (w/v) Fast Green (AppliChem).

### DiI tracing and quantification of dI1 axon guidance at a choice point

DiI tracing of dI1 commissural axons was performed as previously described in detail (Wilson and Stoeckli, 2012). In brief, spinal cords were dissected from the trunk of HH25-26 embryos as an open-book and fixed for 20-60 min at room temperature with 4% PFA in PBS and washed once with PBS. Fast-DiI (5 mg/ml in ethanol; Molecular Probes) was focally injected into the dorsal-most population of commissural neurons (dI1 population). The labeled axons at the midline were visualized using confocal microscopy (Olympus DSU coupled with a BX61 microscope). Only DiI injection sites located in the appropriate dorsal region of the spinal cord, within the level expressing GFP in the FP, were included in the analysis. Because it was not feasible to count axons at individual injection sites, the percentage of axons exhibiting abnormalities was estimated. An injection site was classified as displaying a ’FP stalling’ phenotype if more than 50% of the axons stalled within the FP, or as a ’no turning’ phenotype if more than 50% of the axons that reached the contralateral FP border failed to turn correctly into the longitudinal axis. In some cases, multiple types of phenotypic errors could be observed at a single DiI injection site. The total number of DiI sites for each condition was pooled, and the percentage of normal injection sites was statistically compared across conditions. At least 6 embryos were examined in each condition by a person unaware of the experimental condition.

### Statistical analyses and figure assembly

Statistical analyses were performed with the GraphPad Prism 7.02 software. All data were tested for normality (normal distribution) using the D’Agostino and Pearson omnibus K2 normality test and visual assessment of the normal quantile-quantile plot before choosing an appropriate (parametric or non-parametric) statistical test. For details on data and statistics, see Table S2 Source data. CorelDRAW 2023 version 24.5.0.731 (Corel) was used to create the figures.

## Acknowledgements

We thank Paul Luzio, Michael Davidson, James Hurley, Alissa Weaver, William Gahl and Wesley Sundquist for constructs. We are grateful to Beat Kunz and Tiziana Flego for excellent technical assistance.

## Author contributions

Conceptualization: A.D., E.T.S.; Methodology: A.D.; Validation: A.D., J.M.R.W.; Formal analysis: A.D., J.M.R.W., E.T.S.; Investigation: A.D., J.M.R.W.; Resources: E.T.S.; Data curation: A.D., J.M.R.W.., E.T.S.; Writing - original draft: A.D, J.M.R.W.; Writing - review & editing: A.D., J.M.R.W., E.T.S.; Visualization: A.D., J.M.R.W.; Supervision: E.T.S., A.D.; Project administration: E.T.S.; Funding acquisition: E.T.S.

## Data availability

All relevant data can be found within the article and its supplementary information.

## Supplementary Data

**Movie 1. Nexinhib-20-mediated inhibition of Rab27a induced aberrant dI1 axon guidance at the contralateral floor-plate border.**

Twenty-four hours of ex vivo time-lapse recordings of the ventral midline of spinal cords show Math1::tdTomato-F-positive dI1 axons (black) crossing the ventral midline (dashed green line) of DMSO-treated control or Nexinhib-20-treated samples. dI1 axons turned rostrally in an organized manner in the controls, but many of them showed aberrant trajectories in the presence of Nexinhib20. Maximum projections of z stacks taken every 15 minutes. r, rostral; c, caudal.

---

**Figure S1.**
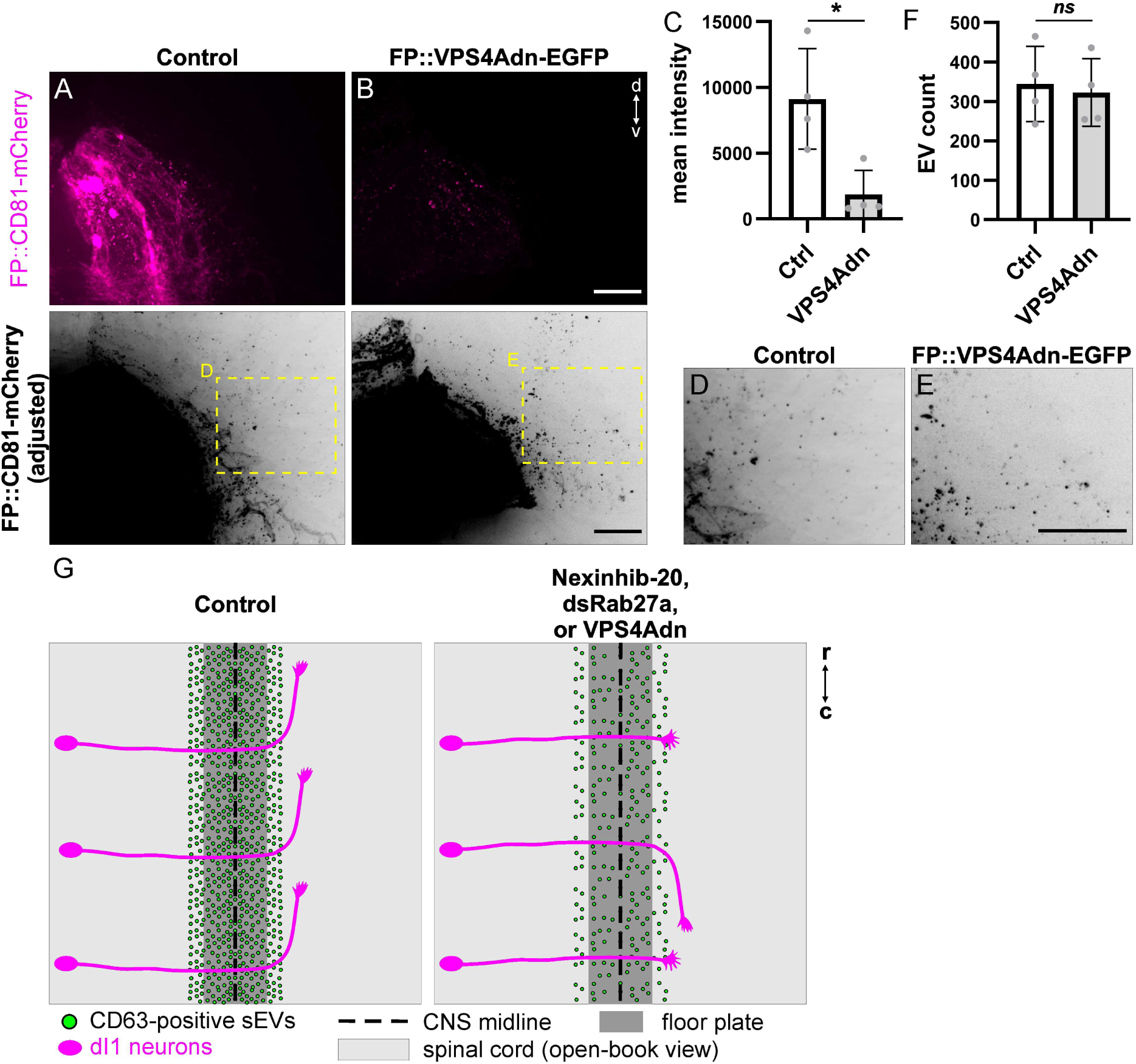
Expression of dominant-negative VPS4 in the floor plate reduced the expression of CD81-mCherry, but not the secretion of CD81-mCherry-positive particles. (A,B) Images of HH23 spinal cord transverse slices showing CD81-mCherry expression in the FP with or without co-expression of VPS4dn. (C) Quantification of the average mean intensity of the CD81-mCherry signal (measured per embryo) within the transfected FP in vivo. Two-tailed unpaired t test. (D,E) Close-up of adjusted images showing secreted CD81-mCherry-positive particles next to the FP. (F) Quantification of the average CD81-mCherry-positive EV count next to the FP. N(embryos)= 4(Control), 4(VPS4dn). Two-tailed unpaired t test. (G) Schematic summary of commissural axon guidance at the FP under control and sEV-depleted conditions. Three different experimental approaches impairing sEV biogenesis and secretion yielded similar post-crossing axon guidance phenotypes. Error bars represent s.d.; d, dorsal; v, ventral; r, rostral; c, caudal; CNS, central nervous system. Scale bars: 20 µm.

**Table S1.** Plasmids used in this study.

| Plasmids |  |  |
| --- | --- | --- |
| plasmid name | concentration used for electroporation | source plasmid and/or reference |
| β-actin::hrGFP<sup>II</sup> | 25 ng/μl | (Wilson and Stoeckli, 2011) |
| Math1::tdTomato-F | 700 ng/μl | (Wilson and Stoeckli, 2011) |
| Hoxa1::CD63-EGFP (CD63-pEGFP C2)* | 6200 ng/μl | Addgene #62964, gift from Paul Luzio, RRID:Addgene_62964 |
| Hoxa1::CD81-mCherry (mCherry-CD81-10)* | 4800 ng/μl | Addgene #55012, gift from Michael Davidson, RRID:Addgene_55012 |
| Hoxa1::ALIX-mCherry (mCherry-hALIX)* | 10 μg/μl | Addgene # 21504, gift from James Hurley, (Lee et al., 2008), RRID:Addgene_21504 |
| Hoxa1::tdTomato-F | 1000 ng/μl | (Dumoulin et al., 2021) |
| β-actin::pHluo-CD63-mScarlet (pHluorin_M153R-CD63-mScarlet)* | 600 ng/μl | Addgene # 172117, gift from Alissa Weaver, (Sung et al., 2020), RRID:Addgene_172117 |
| Hoxa1:hRab27a-EGFP* | 1000 ng/μl | Addgene # 89237, gift from William Gahl, (Westbroek et al., 2008), RRID:Addgene_89237 |
| Hoxa1::VPS4Adn(VSP4-E228Q)* | 1200 ng/μl | Addgene # 80351, gift from Wesley Sundquist, (Votteler et al., 2016), RRID:Addgene_80351 |

